# Enabling reproducible and reusable genetic demultiplexing benchmarking with Nextflow and Apptainer

**DOI:** 10.1101/2025.08.06.668897

**Authors:** Michael P. Lynch, Laurent Gatto, Aedin C. Culhane

## Abstract

Reproducibility, reusability, and portability were identified as key requirements for analysis pipelines in genomics. A previous study identified that less than 4% of notebooks used for biomedical research were fully reproducible. Reusable and portable software are required to ensure compatibility with federated and cloud computing. The field requires computational workflows that meet high standards for reproducibility, reusability, and portability to allow benchmarking to be repeated and updated periodically. This is of particular importance in areas where new methods are frequently released, such as demultiplexing scRNA-seq, a critical early step in most scRNA-seq analysis pipelines.

To address this, we developed *demux_bench* for benchmarking genetic demultiplexing methods in single-cell RNA sequencing, which meets the gold standard for reproducibility of computational workflows, and incorporates best practices for reusability and portability. We used workflow manager Nextflow to enable the simulation and testing of various benchmarking scenarios and methods in parallel from a single pipeline execution. Different experimental configurations can be simulated, including % doublets and class size imbalance. The pipeline includes genotype-free methods, Vireo and souporcell, and additional existing or novel methods can be added modularly. *Demux_bench* is configurable to reproduce specific analyses and generalisable to address new research questions. Software dependencies are handled through containers via Apptainer, allowing portability to different compute environments and avoiding the need for manual installation of software. *Demux_bench* is available on WorkflowHub (https://workflowhub.eu/workflows/1769), and can also be run on Galaxy and other platforms through RO-Crates.

*Demux_bench* facilitates gold standard benchmarking for genetic demultiplexing through reproducibility, reusability, scalability, and portability.

## 1 Introduction

Benchmarking studies underpin bioinformatics analysis by informing method selection based on performance characterisation and highlighting optimal use-cases and limitations. However, challenges remain in performing effective benchmarking studies. Large numbers of tools are published for specific tasks, including demultiplexing, each with different dependencies, including R (Wong et al., 2023), Singularity (Heaton et al., 2020), Python (Xu et al., 2019), conda (Huang et al., 2019), and additional system dependencies. Manual software installation and dependency resolution for many tools can be time-consuming. In the absence of objective measures of performance, methods may be chosen based on other factors such as usability. Additionally, due to the continuous development of new methods, benchmarking studies quickly become outdated and their value diminishes over time (Mangul et al., 2019). Solutions that enable reproducible and reusable benchmarking on multiple platforms are required.

Reproducibility is a challenge in scientific research, including bioinformatics. Issues of reproducibility in data analyses in the life sciences are well documented (Ioannidis et al., 2009; Samuel & Mietchen, 2024). Ioannidis et al. (2009) attempted to reproduce a figure from 18 microarray gene expression studies with two teams of analysts, from which only two figures could be reproduced in principle, and ten could not be reproduced at all. More recently, Samuel and Mietchen (2024) systematically assessed the methods reproducibility (defined as providing sufficient details from the study so that the procedures and data could be repeated exactly) of Jupyter notebooks associated with 3,467 publications indexed in the biomedical literature repository PubMed Central, which mentioned GitHub and Jupyter, where full text was available. These publications were associated with 27,271 Jupyter notebooks that were hosted in 2,660 GitHub repositories. Concerningly, 122 GitHub repositories mentioned were inaccessible. The accessible repositories contained 22,578 Jupyter notebooks written in Python, of which 15,817 notebooks had their dependencies declared in a standard format and 10,388 could have all of their dependencies installed successfully. Of these, only 1,203 notebooks re-ran without any errors, and only 879 produced results identical to those reported in the original notebook. The authors concluded that the majority of published notebooks could not be executed automatically, mostly due to issues related to dependencies, code, or data.

Computational fields are uniquely placed in having the tools to create system-agnostic, verifiably reproducible research, and this applies equally to benchmarking studies as well as primary analyses. Standards for computational reproducibility have been proposed, ranging from a minimum standard of publishing all models, data, and code used in the analysis (bronze), to reproducing the entire analysis with a single command (gold) (Heil et al., 2021). However, implementation and enforcement of such standards vary.

Reusability of benchmarking pipelines extends publications’ value past the original publication. In areas such as demultiplexing scRNA-seq, where new methods are being continuously developed, a static benchmark falls short of the potential value and community need that such studies could provide. Others have proposed models for continuous or live benchmarking, which provide a dynamic account of the field in question (Mallona, Luetge, et al., 2024; Mallona, Soneson, et al., 2024; Soneson & Robinson, 2016), the feasibility of which is dependent on the type of data and methods being benchmarked. FAIR (findable, accessible, interoperable, reusable) (Barker et al., 2022) and similar principles (Freire & Chirigati, 2018) promote reusability and reproducibility of research software, among other attributes. Such standards are already applied to primary analysis tools such as those hosted by Bioconductor (Gentleman et al., 2004; Huber et al., 2015) and applying them to benchmarking pipelines could significantly increase the value of such tools to the community over time.

Due to the diverse range of computing infrastructure and environments, modern bioinformatics tools must be portable to be reusable and reproducible. Modern workflow managers such as Nextflow (Di Tommaso et al., 2017) and Snakemake (Köster & Rahmann, 2012) can be configured to run on multiple environments, including local workstations, high-performance computers (HPC), or cloud environments, with little adaptation to the pipeline configuration, allowing standardised and portable task execution.

Rapid development and updating of methods make tracking and replicating compute environments with specific software versions challenging. Such procedures are inconvenient for local or HPC use but are incompatible with cloud or federated computing applications. Software container systems such as Docker/Podman (Merkel, 2014) and Apptainer/Singularity (Kurtzer et al., 2017) provide a solution by virtualising a specific operating system and software so execution runs independently from the underlying operating system, and these can be integrated into existing modern workflow managers.

Identifying dataset properties which are predictive of method performance (Strobl & Leisch, 2024) requires the ability to parameterise and parallelise benchmarking studies across scenarios and fully utilise compute resources with minimal manual intervention. This is well within the capabilities of modern workflow managers, but requires additional investment of time when designing workflows.

## 2 Statement of need

Reproducibility, reusability, and portability were identified as key features of analysis pipelines in genomics (Ahmed et al., 2021; Wratten et al., 2021). With these attributes in mind, we extended existing genetic demultiplexing simulation and benchmark techniques (Weber et al., 2021) into a reproducible and reusable tool *demux_bench*, which systematically tests against experimental parameters and is configurable to reproduce specific analyses or generalise to address new research questions.

Multiplexing, which is the simultaneous sequencing of multiple biological samples in the same sequencing lane, and subsequent successful computational demultiplexing of scRNA-seq, is a crucial early step in scRNA-seq that increases the scale of experiments while increasing cost effectiveness. Optimal computational demultiplexing is required to ensure correct cell assignment and prevent loss of valuable data. Typical data loss of 1-17% of cells (M. P. Lynch et al., 2024) is common with losses as high as 50% observed (Howitt et al., 2023). Misassignment of cells can produce substantial errors that can render experimental results invalid.

The growing number of available demultiplexing algorithms (Heaton et al., 2020; Huang et al., 2019; Wong et al., 2023; Xu et al., 2019) requires systematic, reproducible, and reusable benchmarking solutions to properly evaluate performance and address the asynchronous gap between methods development and independent benchmarking currently observed (Figure 1A). Previous benchmarking studies (Weber et al., 2021) developed *in silico* SNP simulation methods with reliable ground truth to determine if genetic demultiplexing was suitable for cancer samples. However, only Vireo (genotype-free) (Huang et al., 2019) and Demuxlet (known-genotype) (Kang et al., 2018) methods were tested, and so genotype-free demultiplexing methods remain without a comprehensive independent benchmark. Hashing-based demultiplexing methods have been more frequently benchmarked and have better coverage in independent benchmark studies (Figure 1B).

**Figure 1.**
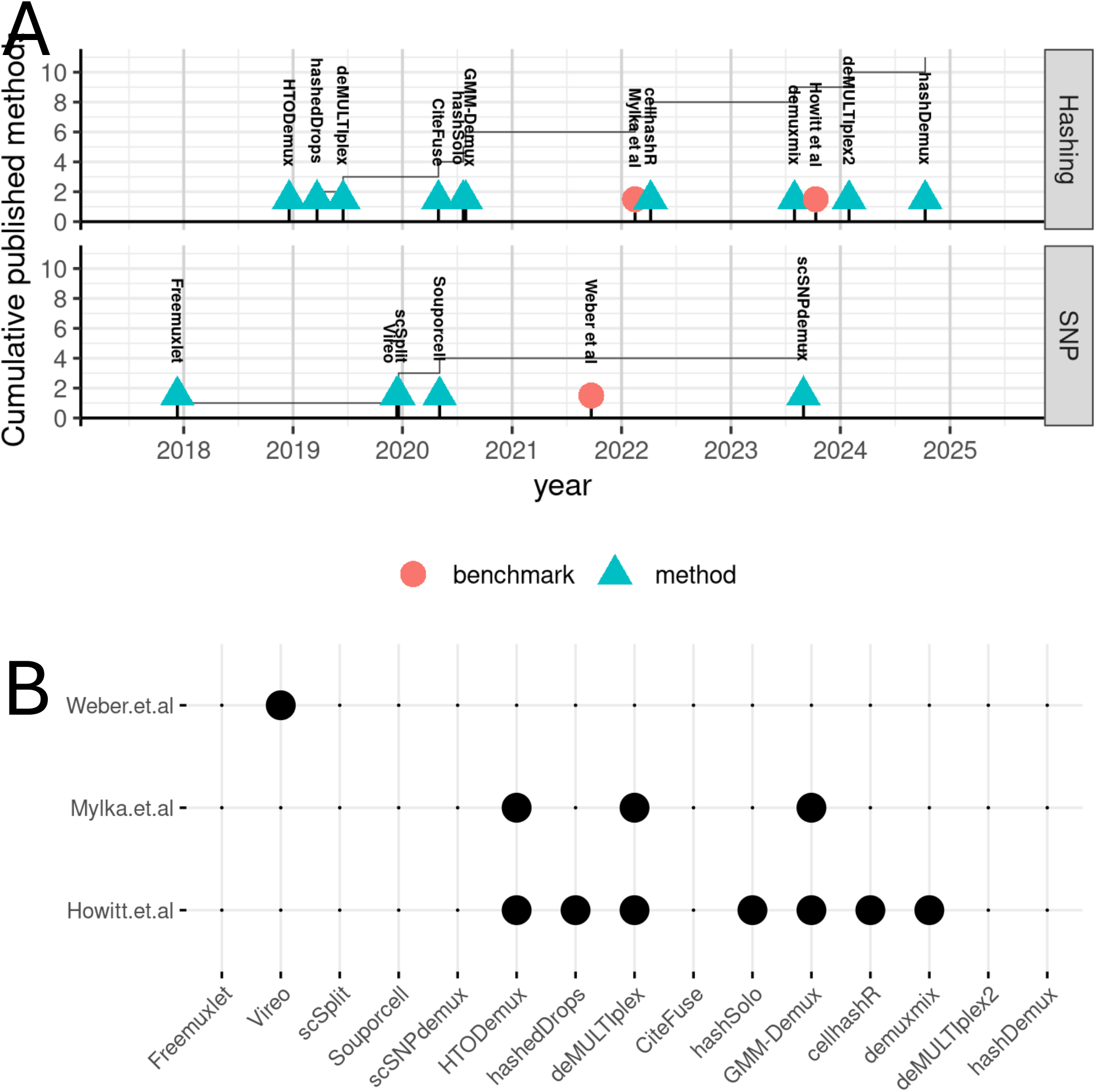
Coverage of hashing and genetic demultiplexing methods in independent benchmarking studies. A) Comparison of publication timelines for methods vs. independent benchmark studies for hashing and genetic demultiplexing.^1^B) Coverage of hashing and genetic demultiplexing methods in independent benchmarking studies.

We describe a Nextflow tool, *demux_bench*, for data simulation and benchmarking of genetic demultiplexing methods. Raw data and software containers are fetched as part of the pipeline, reducing the required dependencies to Nextflow (Di Tommaso et al., 2017) and Apptainer (Kurtzer et al., 2017). Nextflow allows the pipeline to run in a relatively system-agnostic manner, compatible with local workstations, HPCs, and cloud-computing platforms. The workflow meets the gold standard for reproducibility in the life sciences (Heil et al., 2021) and is executable on an example dataset (10X Genomics, 2021) with a single command. It is also configurable to generalise to different datasets and simulation scenarios, and is available on WorkflowHub (https://workflowhub.eu/workflows/1769) and GitHub (https://github.com/michaelplynch/demux_pipeline).

## 3 Methods and Results

*demux_bench* is implemented in Nextflow (Di Tommaso et al., 2017) due to its emphasis on scalable and reproducible data-intensive workflows. Nextflow uses a dataflow programming model, which uses asynchronous queues (channels) to pass data between different computational tasks (processes). The flow of information can be visualised as a directed acyclic graph. The pipeline contains one optional preprocessing section for fetching example raw data (not shown) and two core sections for data simulation and methods testing (Figure 2).

**Figure 2.**
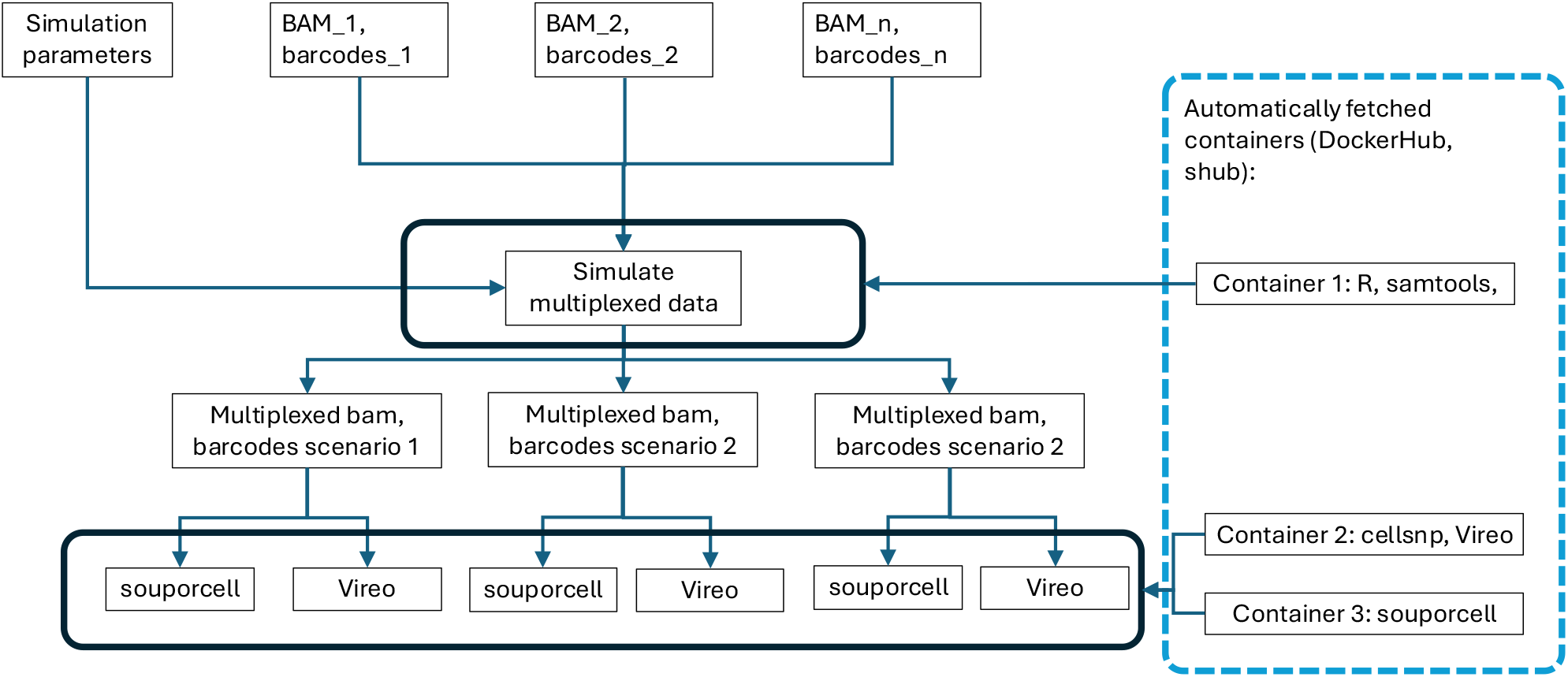
Directed acyclic graph describing core sections of demultiplexing simulation and benchmarking workflow. Doublet simulation process is expanded in Figure 3. Workflow dependencies for doublet simulation, Vireo, and souporcell method are containerised and automatically fetched from standard repositories (DockerHub, shub).

### 3.1 Inputs

#### 3.1.1 Raw data

Input raw data consists of per-sample BAM and barcodes files and is defined by one of two modes, set by a flag defined in nextflow.config, which determines how the input data channels are created. The default mode (download flag=false) is to specify file paths to all inputs (BAM files, barcodes, reference genome and variants) in the nextflow.config file (Table 1). The pipeline is designed to accept an arbitrary number of input samples (BAM files and corresponding barcodes). The non-default mode (download flag=true) runs an initial process to download an example dataset from 10X Genomics (10X Genomics, 2021) to demonstrate gold standard reproducibility. The output of this process is two channels containing file paths for BAM and barcodes files, respectively. This initial step can be run by setting ‘—download flag true’ when executing the pipeline on the command line and only needs to be run once to prevent unnecessary network traffic. After the initial download, data is stored in the project directory in ‘data/input’ and can be referenced in nextflow.config for subsequent executions of the pipeline.

**Table 1:** Benchmarking pipeline input data.

| Parameter | Description |
| --- | --- |
| bam_path | Paths to per-sample bam files |
| barcodes_path | Paths to per-sample barcodes files, the order of samples should correspond to bam_path. Barcodes should be in .tsv format with one column and no column name. |
| ref | Path to 10X Cell Ranger-compatible reference genome |
| common_variants | Path to common variants (.vcf) |

The corresponding reference genome and variants file must be manually downloaded and specified in the nextflow.config file.

*demux_bench* accommodates 10X Genomics single-cell sequencing data, and therefore expects RNA-sequencing data that were prepared using the 10X Genomics Cell Ranger count pipeline (Zheng et al., 2017). Data output from the Cell Ranger multi pipeline is compatible with slight modifications to the barcodes file. A database of common variants, adapted from Huang et al. (2019), is provided (M. Lynch, 2025) but can be substituted with a user-provided .vcf (variant call format) file. These are stored within the project directory for use by downstream processes.

#### 3.1.2 Benchmarking parameters

Benchmarking scenarios are defined in params.csv, with one row per benchmarking scenario (Table 2). The first column represents the unique key to identify that scenario in the form of a string. The second column specifies the percentage of doublets simulated. The remaining columns determine the number of cells to subsample from each input sample based on the order in which they were defined in bam path and barcodes path.

**Table 2:** Example specification of simulation parameters.

| key | pcdoub | n_1 | n_1 | .. | n_i |
| --- | --- | --- | --- | --- | --- |
| a | 5 | 5,000 | 5,000 | .. | 5,000 |
| .. | .. | .. | .. | .. | .. |
| j | 50 | 5,000 | 5,000 | ... | 5,000 |

*demux_bench* requires a matching key to be specified for each simulation scenario to ensure correct and efficient processing of data. In Nextflow, channels are defined as a value (which can be consumed multiple times, e.g., path to reference genome) or a queue, where each element of the queue is supplied to a process once (e.g., parsing individual BAM files), after which the channel is depleted. Queue channels, which are produced from the output of a process, emit elements in the order in which the sub-processes complete. Importantly, this does not necessarily match the order of the input to said process. To address this, matching keys are defined using tuples, where the first element of the tuple is a unique string identifying the benchmarking scenario being simulated. Channel operators can then join channels (e.g., barcodes and corresponding BAM files) based on this string to guarantee correct matching of elements from different channels.

#### 3.1.3 Software

*demux_bench* requires Apptainer to fetch and execute processes through containers. The software containers created for this pipeline are hosted on DockerHub (https://hub.docker.com/repository/docker/mplynch28/demux_sim & https://hub.docker.com/repository/docker/mplynch28/demux_vireo). Soupor-cell is distributed as a Singularity container on shub. Nextflow downloads and caches these containers automatically upon execution of the respective process.

### 3.2 Data simulation

The second section of the pipeline simulates *in silico* SNP data representing different benchmarking scenarios using the raw sequencing data as input. *demux_bench* enables the simultaneous execution of different data simulation scenarios and subsequent testing of different methods (Heaton et al., 2020; Huang et al., 2019) on them.

The core processing steps of the pipeline are briefly outlined below (Table 3). Processes are only simulated as often as they need to, e.g., the BAM files are parsed and merged only once, regardless of how many scenarios are being evaluated (Figure 3) to avoid unnecessary computation and allocation of compute resources.

**Table 3:** Description of workflow processes for simulating doublets.

| Process | Function |
| --- | --- |
| PARSE | Sample-specific suffixes are added to barcodes in each BAM file. |
| MERGE | Per-sample BAM files are merged into one ‘multiplexed’ BAM file. |
| RENAME_BCS | Sample-specific suffixes are added to barcodes in each barcodes file. |
| SUBSET_BCS | Barcodes are subsampled based on input from params.csv. |
| MERGE_BCS | Per-sample barcodes files are merged into one ‘multiplexed’ file. |
| GENERATE_LOOKUP | Barcodes are randomly selected to be renamed to another randomly selected barcode to represent the desired % doublets. |
| SIM_DOUBS | Barcodes are renamed as per the generated lookup. |

**Figure 3.**
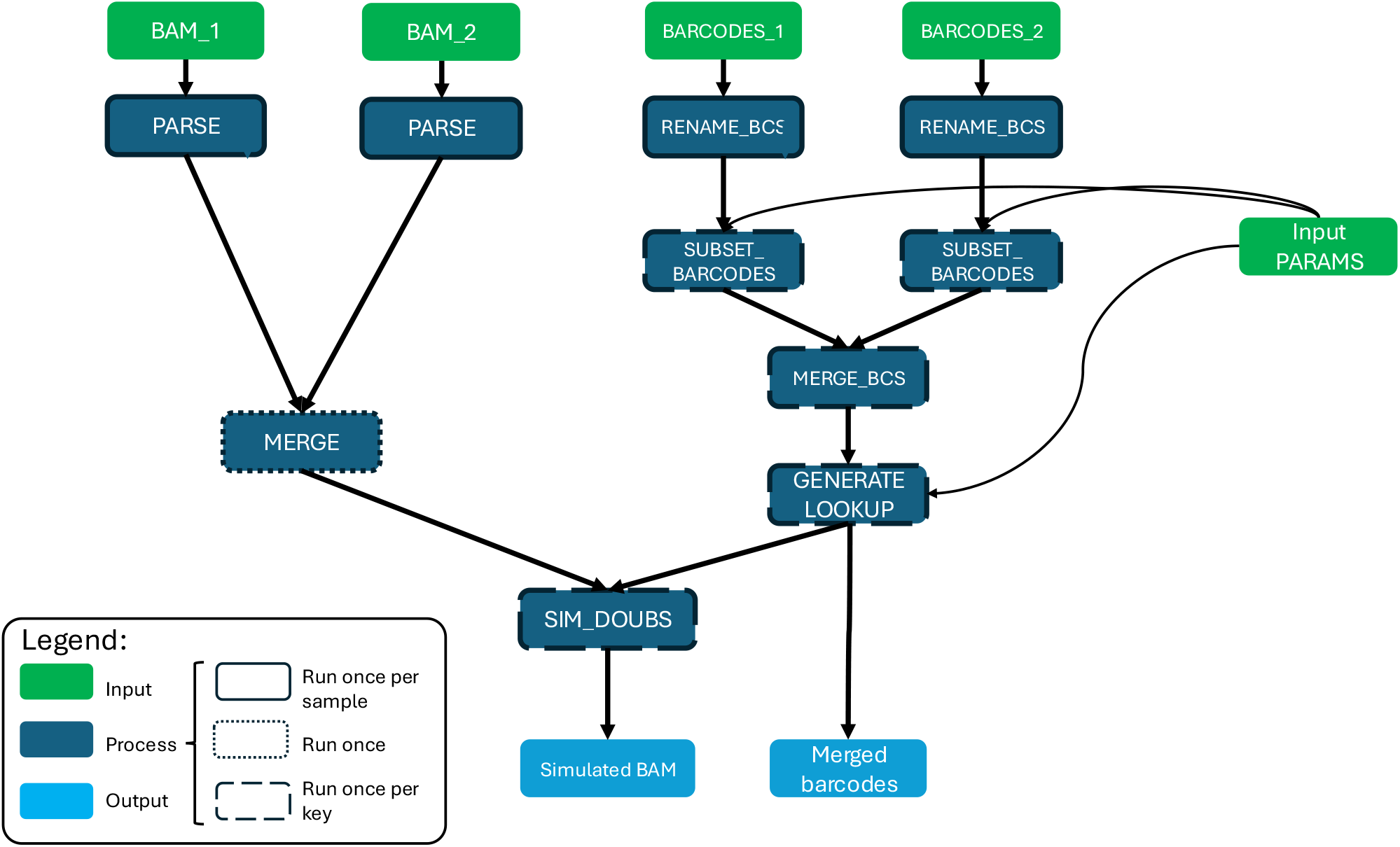
Directed acyclic graph detailing doublet simulation workflow. Data are simulated based on scenarios described in a configuration .csv file. Workflow logic avoids the unnecessary rerunning of pipeline steps.

### 3.3 Features

#### Reproducible

After downloading input data and adjusting configuration for the environment, *de-mux_bench* can be rerun with a single command, meeting the gold standard for computational reproducibility in the life sciences (Heil et al., 2021).

#### Reusable

*Demux_bench* can be configured to run on different datasets with arbitrary numbers of samples to be multiplexed.

#### Scalable

*Demux_bench* simulates data for benchmarking scenarios and runs methods in parallel. Our implementation allows methods to be added in a modular fashion. Data can be simulated for various scenarios based on % doublets and number of cells per sample and are simulated and evaluated in parallel (Figures 2,3,4).

**Figure 4.**
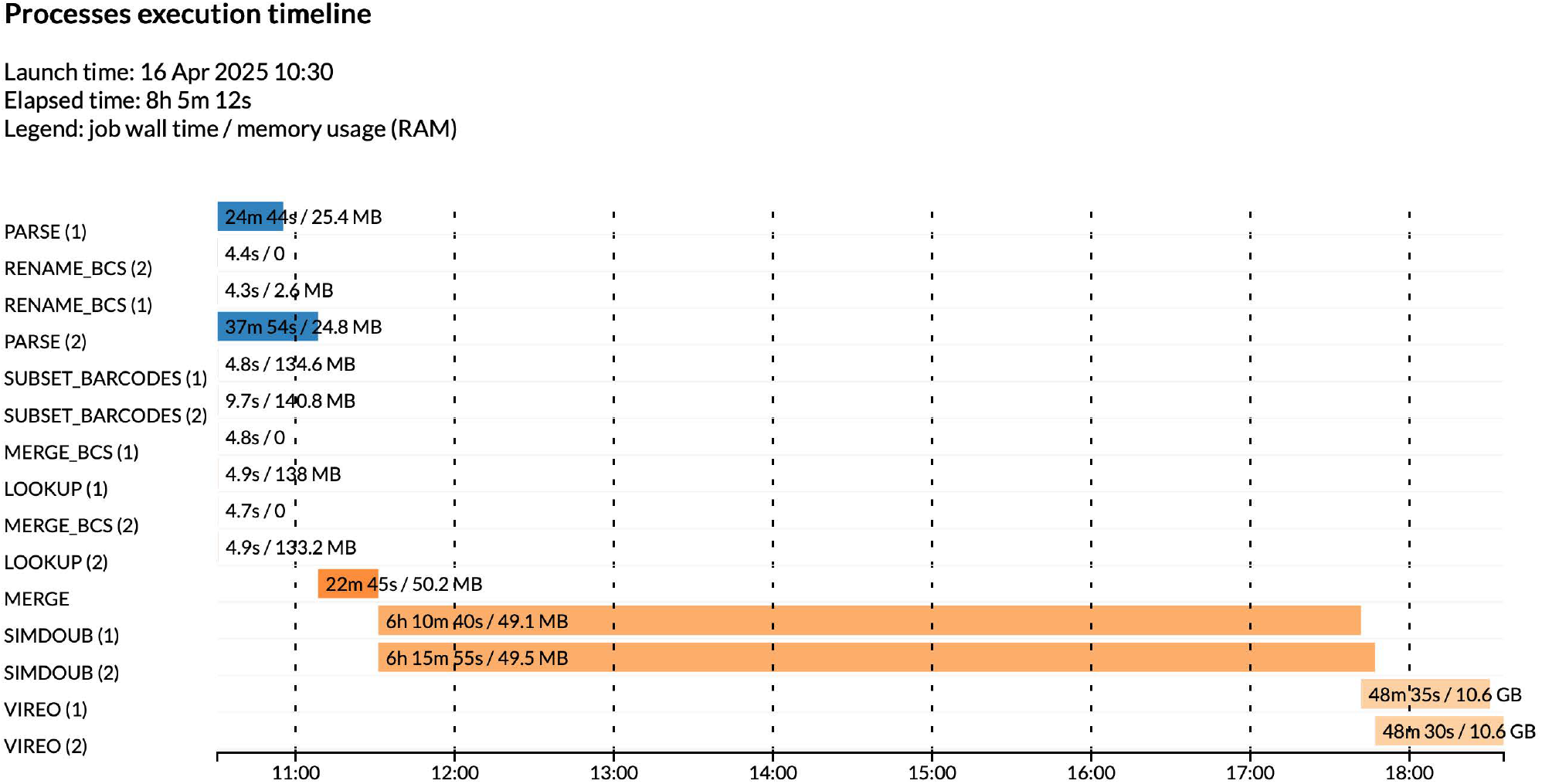
Process execution timeline for sample 10X Genomics cell line dataset. Each bar represents a process that may be run more than once, depending on inputs. Different tasks from the same process are coloured the same. The coloured section of the bar represents real execution time. Figures within each bar are wall time and memory usage (RAM).

Fully containerised: All required software and system dependencies are packaged into containers available on DockerHub (Figure 2) (https://hub.docker.com/repository/docker/mplynch28/demux sim and https://hub.docker.com/repository/docker/mplynch28/demux vireo for doublet simulation and Vireo Docker containers, respectively). Souporcell is already distributed through a Singularity container.

### 3.4 Configuration

Configuration parameters for process execution are described in Table 4. The execution time of different processes varies between hours for processing BAM files and seconds for processing barcodes files (Figure 4, Intel Xeon Gold 6342 CPU @ 2.80GHz).

**Table 4:** Pipeline execution configuration parameters.

| Parameter | Default value | Description |
| --- | --- | --- |
| process.executor | 'slurm' | Job scheduler |
| apptainer.enabled | true | Run through Apptainer containers |
| process.module | 'apptainer' | Whether Apptainer needs to be loaded thorough modules environment |

For workload managers such as SLURM, each subprocess is submitted as a separate job. Further configuration of resource allocation is possible with Nextflow and may be desirable depending on the compute environment. Additional parameters for configuration can be added to nextflow.config.

### 3.5 Installation

*demux_bench* is available on WorkflowHub (https://workflowhub.eu/workflows/1769) or the repository can be cloned from GitHub (https://github.com/michaelplynch/demux_pipeline), and requires Nextflow and Apptainer to run. All remaining dependencies are handled through containers (Figure 2). The pipeline can then be run on example data using

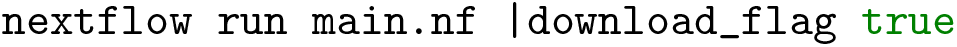

from the top level of the project directory. The associated RO-Crate, available through WorkflowHub.eu, can be imported into, and executed on, Galaxy.

## 4 Discussion

The community continues to raise the standard for describing computational work for primary analyses and benchmarking methods alike. Benchmarking studies performed on realistic data that generate generalisable findings are key for understanding the performance and limitations of methods developed to analyse scRNA-seq data, amongst other high-throughput assays.

We developed an open-source pipeline, *demux_bench*, for the simulation and testing of genotype-free genetic demultiplexing methods across different scenarios for % doublets and number of cells per sample that meets the gold standard for computational reproducibility. The pipeline uses software containerisation through Apptainer (Kurtzer et al., 2017) and DockerHub, and workflow manager Nextflow (Di Tommaso et al., 2017) to ensure reproducibility, reusability, scalability, and portability.

*Demux_bench* may be reused for future independent benchmarking or method development.

The pipeline is an important development in evaluating the performance of genotype-free genetic demultiplexing methods. It addresses testing of different scenarios which can be parameterised and simulated and were found to impact method performance, such as class imbalance and % doublets (M. P. Lynch et al., 2024). However, this is not an exhaustive set of potentially impactful factors, and testing of other unforeseen factors may require some adaptation of the pipeline.

Additionally, we focused on genotype-free genetic demultiplexing methods, a subset of the available genetic demultiplexing methods. Other methods require per-sample genotypes to be known *a priori*, which is not incorporated into the current workflow. The release of multi-modal and hybrid methods will require more complex pipelines to handle the simulation of different modalities of data in parallel. Reproducible benchmarking pipelines require persistence in software dependencies and input data.

For software, we addressed handling changes in versioning and maintenance through container technologies. For raw data, many specific-purpose repositories (GEO, EGA) exist, which are likely to provide persistent access to data. However, as we move towards greater protection of patient privacy by storing genomics data in controlled-access databases such as dbGaP, containerised and portable pipelines will be crucial for analysing private data. Secondary files such as reference genomes, formatted for use with 10X Genomics software, are currently not guaranteed to persist.

Live benchmarking was proposed by Soneson and Robinson (2016), who created a package for standardised benchmarks for genomic tasks. Methods could be executed live and results visualised through an R Shiny interface. However, nine years later, many studies still fail to meet this level of transparency. Their recent publication (Mallona, Soneson, et al., 2024) offers perspectives on a wish list for bioinformatics benchmarking and extends the concept of live benchmarking to continuous benchmarking, where benchmarking systems are standardised and thus can be contributed to or corrected as new methods or versions become available. Containerised pipelines such as this underpin live benchmarking processes in applications such as genetic demultiplexing that require complex simulation processes and expensive computation. For an application such as hashing demultiplexing (Howitt et al., 2023), which is less data and compute-intensive, continuous or live systems provide a solution to assessing method performance across many datasets relatively cheaply. This would extend current available benchmarking in this area (Howitt et al., 2023; Mylka et al., 2022) to understanding which methods are performant in different experimental scenarios or dependent on specific experimental parameters (Strobl & Leisch, 2024).

As the field increasingly relies on federated and cloud computing, features such as reproducibility, reusability and portability will be essential for software tools and pipelines. Compatibility with RO-Crates (Zimmerman & Langefeld, 2021) through WorkflowHub allows for interoperability with Galaxy and other workflow managers.

## 5 Conclusion

*demux_bench* allows for minimal effort or intervention to reproduce and reuse genotype-free genetic demultiplexing benchmarking by design. Software dependencies are containerised, and the relevant containers and data (configuration dependent) are fetched on execution of the pipeline. Modular Nextflow processes make the pipeline readily extendable, facilitating continuous benchmarking to new methods.

## Footnotes

1 Date taken as publication date of associated manuscript except for Freemuxlet, which in the absence of a publication is taken as the publication date of Demuxlet, which are packaged together.

